# Effect of Transcranial Light Stimulation on the Neurovascular Unit in the Human Brain

**DOI:** 10.1101/2025.04.23.649937

**Authors:** Chenguang Zhao, Zhilin Li, Zhaohuan Ding, Keyao Zhang, Yang Li, Long Zhang, Qihong Zou, Zhibing Gao, Zaixu Cui, Xiaoli Li

**Affiliations:** Chinese Institute for Brain Research, Beijing 102206, China & Peking Union Medical College, Beijing, 102206, China; Shien-Ming Wu School of Intelligent Engineering, South China University of Technology, Guangzhou, 510640, China; School of Biomedical Engineering, Faculty of Medicine, Dalian University of Technology, Dalian, 116024, China; School of Artificial Intelligence, Beijing University of Posts and Telecommunications, Beijing, 100876, China; McGovern Institute for Brain Research, Peking University, 5 Yiheyuan Road, Haidian District, Beijing, China; National Center for Mental Health, Beijing, 100029, China; Pazhou Laboratory (Guangzhou) & School of Automation Science and Engineering, South China University of Technology, Guangzhou 510641, China

**Author notes:** Corresponding Author : Chenguang Zhao; Zaixu Cui; XiaoLi Li. These authors contributed equally to this work (Chenguang Zhao, Zhilin Li, Zhaohuan Ding).

**Keywords:** Transcranial light stimulation, Neurovascular unit, TMS-EEG, tLS-fMRI, tLS-EEG

## Abstract

Transcranial light stimulation (tLS) is emerging as a non-invasive approach for enhancing brain function and treating neurological disorders; however, its impact on the human neurovascular unit (NVU) remains poorly understood. Herein, we combined photon transport modeling with multimodal neuroimaging to reveal how light influences vascular and neuronal responses in the human brain. Simulations of photon propagation through transcranial tissue captured key scattering and attenuation patterns, guiding the localization of light effects in vivo. Using simultaneous functional magnetic resonance imaging and arterial spin labeling, we showed that tLS significantly increased blood oxygenation level-dependent signals and cerebral blood flow in the light-affected regions. These hemodynamic changes co-occurred with a reduction in cortical excitability, as revealed by electroencephalographic source reconstruction and transcranial magnetic stimulation-evoked potentials. To probe the underlying mechanism, we incorporated inhibitory neural inputs into the computational NVU model. The model predicted that tLS enhances inhibitory neuronal activity and nitric oxide release, driving vasodilation and elevating metabolic support. These findings revealed that transcranial photons can differentially modulate neuronal and vascular components of the NVU—suppressing excitability while promoting perfusion—thereby suggesting a novel therapeutic avenue for targeting neurovascular dynamics in cognitive and clinical applications.

## Introduction

Near-infrared (NIR) wavelengths (600–1100 nm) can non-invasively penetrate skull tissues (Wu et al., 2022), reaching the cerebral cortex where mitochondrial cytochrome c oxidase (CCO) acts as the primary NIR photoacceptor. This mechanism promotes adenosine triphosphate (ATP) synthesis through photon absorption (Karu et al., 2008). Human studies have shown that non-invasive transcranial light stimulation (tLS) improves cognitive performance (Zhao et al., 2022; Lee et al., 2023) and clinical potential in neurological conditions, including stroke (Zivin et al., 2009; Kim et al., 2022), traumatic brain injury (Longo et al., 2020), Alzheimer’s disease (Tao et al., 2021), pathological fear (Zaizar et al., 2021) and hemorrhage (Li et al., 2023).

Our current understanding of mitochondrial responses to NIR light is mainly limited to observations at the submicron scale. In contrast, at the macroscopic level of the human brain, the smallest observable unit contains numerous mitochondria distributed throughout vascular systems, neurons, and glial cells (Mosharov et al., 2025). Beyond ATP production, mitochondria play critical roles in neurotransmitter release, signaling, and modulation of neuronal excitability (Wang et al., 2024). In animal models, light-induced nitric oxide (NO) release enhances cerebral blood flow (CBF) (Uozumi et al., 2010), whereas human studies have reported localized increases in oxygenated hemoglobin (Wang et al., 2017) and frequency-specific modulation of electroencephalographic (EEG) activity (Li et al., 2024). Growing evidence indicates that the neurovascular unit (NVU) may be a promising target for tLS (Yan et al., 2025). However, due to a methodological and conceptual gap in scaling cellular mechanisms to integrated human brain function, how tLS influences the NVU in the human brain remains unexplored.

The NVU exhibits high conservation in its core functional mechanisms. For example, in mice, NO-mediated vasodilation is driven by specific interneurons, whereas in primates, the relationship between local field potentials and blood oxygenation level-dependent (BOLD) signals more closely resembles that in humans (Schaeffer et al., 2021). The mechanisms of light-induced modulation established in animal models can be preserved and extended to the analysis of primate data, and further propagated to in vivo studies in humans. It is essential to observe and predict the multifaceted influences on neurons and blood vessels as well as their intricate interplay at appropriate spatial and temporal scales. Thus, guided by cross-species NVU models, we employ multimodal neuroimaging techniques—including functional magnetic resonance imaging (fMRI), EEG, and arterial spin labeling (ASL) — to characterize alterations in neural electrophysiology and cerebral hemodynamics. These observed changes correlate with NO-mediated signaling pathways and vascular smooth muscle cell (VSMC)/capillary dynamics, which have been established as key mechanistic processes underlying tLS in previous studies. (Yan et al., 2025).

A key limitation in assessing NVU dynamics is defining the spatial boundaries of photobiological effects throughout human cranial anatomy. We first employed individualized Monte Carlo (MC) simulations based on structural imaging to map intracranial light energy distributions, enabling the precise localization of brain regions exposed to biologically relevant doses of NIR light. The MRI-compatible tLS-fMRI system permitted millimeter-resolution whole-brain imaging while simultaneously quantifying hemodynamic responses to tLS in real time. In the illuminated regions, we measured changes in CBF and cerebral metabolic rate of oxygen (CMRO□) using dual-echo pseudo-continuous ASL (Faraco et al., 2015). To examine the electrophysiological responses of tLS, we reconstructed spontaneous neuronal activity to assess the excitation/inhibition (E/I) balance in the targeted areas (Sohal et al., 2019). Additionally, we employed transcranial magnetic stimulation coupled with EEG (TMS–EEG) to directly probe cortical excitability and assess the dynamic reactivity to tLS.

Our results revealed that tLS suppresses cortical excitability while enhancing CBF and metabolic activity. These effects provide a mechanism linking photon attenuation to GABAergic inhibition and NO-mediated vasodilation in humans. Our findings provide converging evidence of a coordinated photobiological response at the NVU level, offering mechanistic insights that can bridge the gap between cellular models and human brain function. This study also lays the foundation for the rational design of personalized tLS interventions, thereby advancing the field of precision light stimulation.

## Results

### High-Precision MRI-Compatible tLS System for Spatially Registered Neurovascular Unit

We developed a high-precision, MRI-compatible, spatial co-registration tLS system to assess the effects of tLS-induced NVU responses in brain tissues. A high-precision laser system can more accurately stimulate and measure the light path within tissues to reliably analyze the effects of tLS with a narrow laser linewidth and high uniformity degree. A laser source with a central wavelength of 1064.17 nm and a linewidth of 0.064 nm in NIR-II was employed (Fig. 1a) and homogenized to 96.34% using a beam shaper (Fig. 1b) to achieve a high-precision incident light, thereby ensuring that the optical characteristics across the beam profile were consistent and uniform. To ensure that light itself does not interfere with MRI and is not affected by the MRI magnetic field, the light delivery system was constructed using MRI-compatible materials, including a polyvinyl chloride (PVC) protective layer, an acrylate resin coating, and an SiO□ core for the optical fiber. The beam delivery components consisted of an Al□O□ heat sink, a nylon structure, and BK7 glass (Fig. 1c). All components were confirmed by MRI-compatible testing. The generated laser was delivered to the MRI scanning room through two MRI-compatible optical fibers (Extended Data Fig. 1a). Light was transmitted to the MRI magnet alongside the participant using beamformers and an MRI-compatible rubber headgear affixed to the forehead (Extended Data Fig. 1b). The forehead, which is free of hair, is an ideal target area for tLS. We divided it into 10 positions arranged in a two × five grid (Extended Data Fig. 1a-b,d). The spatial positions of these 10 light spots were determined using vitamin E (VE) markers visible in the T1 image (Extended Data Fig. 1c) and constrained by the design of the headgear (Extended Data Fig. 1d; positions shown in Fig. 1e). Two T1 sequences were acquired, one for VE marker localization and the other for tissue segmentation (Extended Data Fig. 1c). These data enabled spatial co-registration of functional signals—CBF, BOLD, and CMRO□—acquired through dual-echo sequences (Fig. 1d). The original T1-weighted MRI image (Fig. 1e) formed the anatomical basis for cortical segmentation using the CAT12 toolbox. The resulting segmented image (Fig. 1e) displayed distinct neuroanatomical regions in a color-coded scheme, which directly supported our photon propagation simulations in heterogeneous brain tissues. The upper-right position of the individualized light stimulation matrix corresponded to the right dorsolateral prefrontal cortex (Extended Data Fig. 1f) as the putative individual site of stimulation. The distribution of individualized target centers is shown for all 43 participants (Fig. 1f). Next, we defined the individualized incidence direction of the beam spot (Extended Data Fig. 1g). The precise and safe delivery of laser light into the MRI scanner, along with the accurate spatial registration of tLS parameters, particularly the target center and normal angle of the incident beam, are critical for establishing the initial conditions required for subsequent light path simulations. Theoretically, this process enhances the precision and efficacy of tLS in brain tissue, leading to more accurate experimental outcomes.

**Fig. 1.**
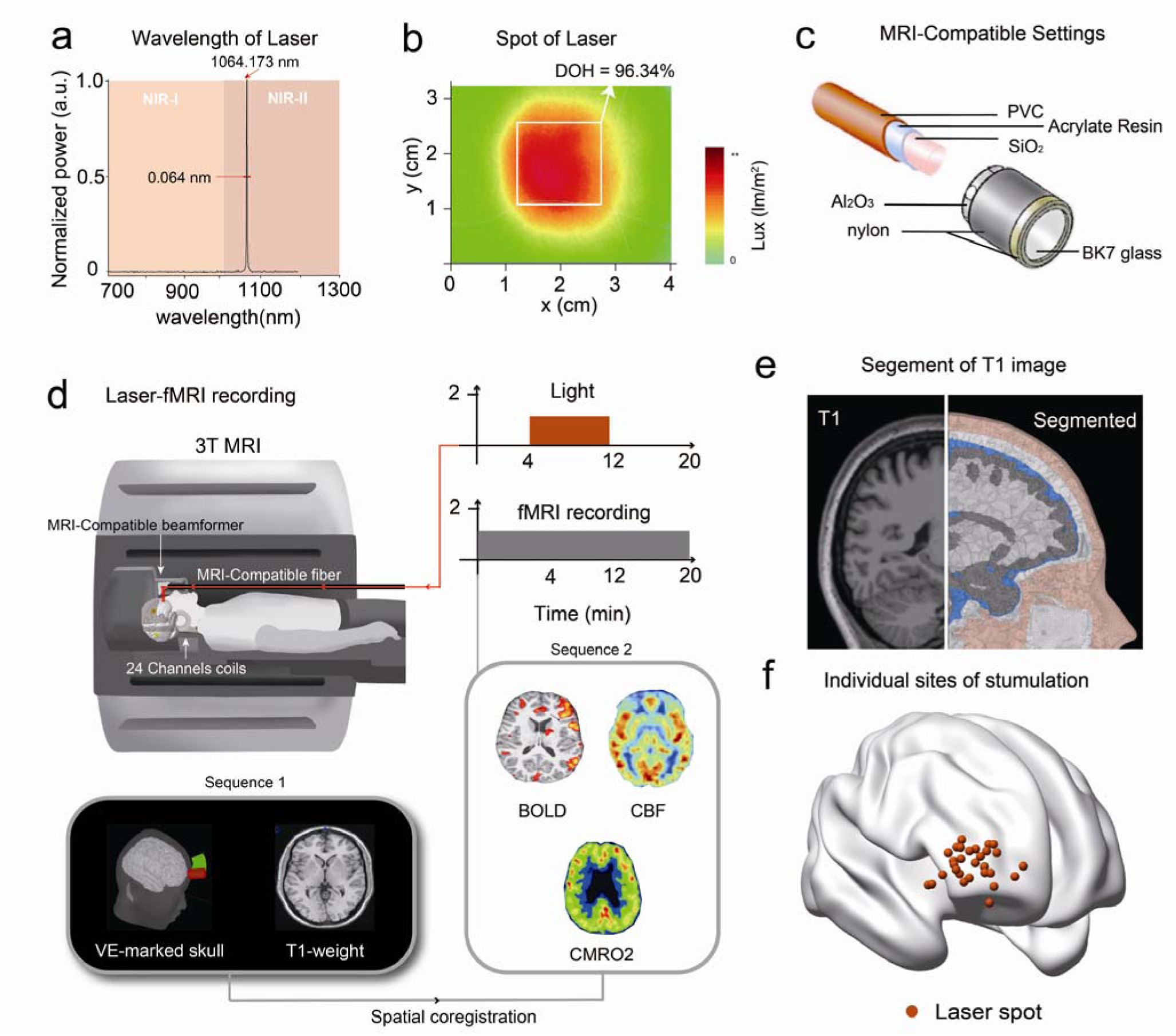
Imaging data recording and co-registration. (**a)** Spectral analysis of the laser shows a central wavelength of 1064.173 nm with a linewidth of 0.064 nm. (**b)** 96.34% of the normalized power falls within the 1064±0.1 nm range, approximating a pure 1064 nm light source. **(c)** The transmission device of light comprises a PVC protective layer, an acrylate resin coating, and an SiO_2_ inner core. The beam-focusing assembly includes an Al_2_O_3_ heat sink, a nylon structural frame, and BK7 glass. All components have passed MRI-compatible testing. **(d)** MRI-compatible tLS recording configuration. The laser is introduced into the main magnet through MRI-compatible optical fibers. A specialized MRI-compatible beam shaper is used to alter the direction and spatial coherence of the laser. Given the use of a 24-channel coil for imaging, the optical device’s design must accommodate the internal space of the coil. MRI recording protocol: simultaneous fMRI-ASL recording was synchronized with tLS, following a 4-min baseline period, 8-min light stimulation, and 8 min post-stimulation. MRI acquisition sequences: T1 image for VE-marked position and structural brain (Sequence 1); Dual-echo sequence for BOLD signals, CBF, and CMRO_2_ (Sequence 2). The former provides spatial information for co-registering the latter. **(e)** T1 structural image (left) were segmented into different brain tissues (right), including skin (pink layer), skull (white layer), cerebrospinal fluid (blue layer), gray matter (gray layer), and white matter (corrugated layer). **(f)** Laser spot for individualized stimulation on cerebral cortex. The red dots indicate the center of each participant’s laser spot projection on the cortex. ASL: arterial spin labeling; BOLD: blood oxygenation level-dependent; CBF: cerebral blood flow; CMRO_2_: cerebral metabolic rate of oxygen; fMRI: functional magnetic resonance imaging; MRI: magnetic resonance imaging; PVC: polyvinyl chloride; tLS: transcranial light stimulation; VE, vitamin E.

### Monte Carlo Simulation of Transcranial Light Propagation and Individualized Attenuation Mapping

Because non-invasive interventions cannot detect the energy attenuation of intracranial photons, we simulated the transcranial propagation of laser light (Fig. 2a-b) using an MC method, which simulates the photon at 1064 nm traveling through the transcranial tissue layers as it reflects or refracts based on light properties (Extended Data, Table 1). By co-registering structural T1 images with individual laser configurations, we can perform personalized photon flux simulations and obtain a transcranial light attenuation map. For each 10-dB attenuation, an individualized region of interest (ROI) was constructed based on simulations of light propagation through the individual’s head (Fig. 2c). The results (Extended Data Fig. 2b) revealed that laser light penetration exhibited exponential attenuation in the brain tissue, characterized by the depth of photon energy along the normal vector at the point of incidence (Extended Data Fig. 2a). This relationship follows the equation y = 0.0004*e^-0.376x^ (*R*^2^=0.986), consistent with the Beer–Lambert law.

**Fig. 2.**
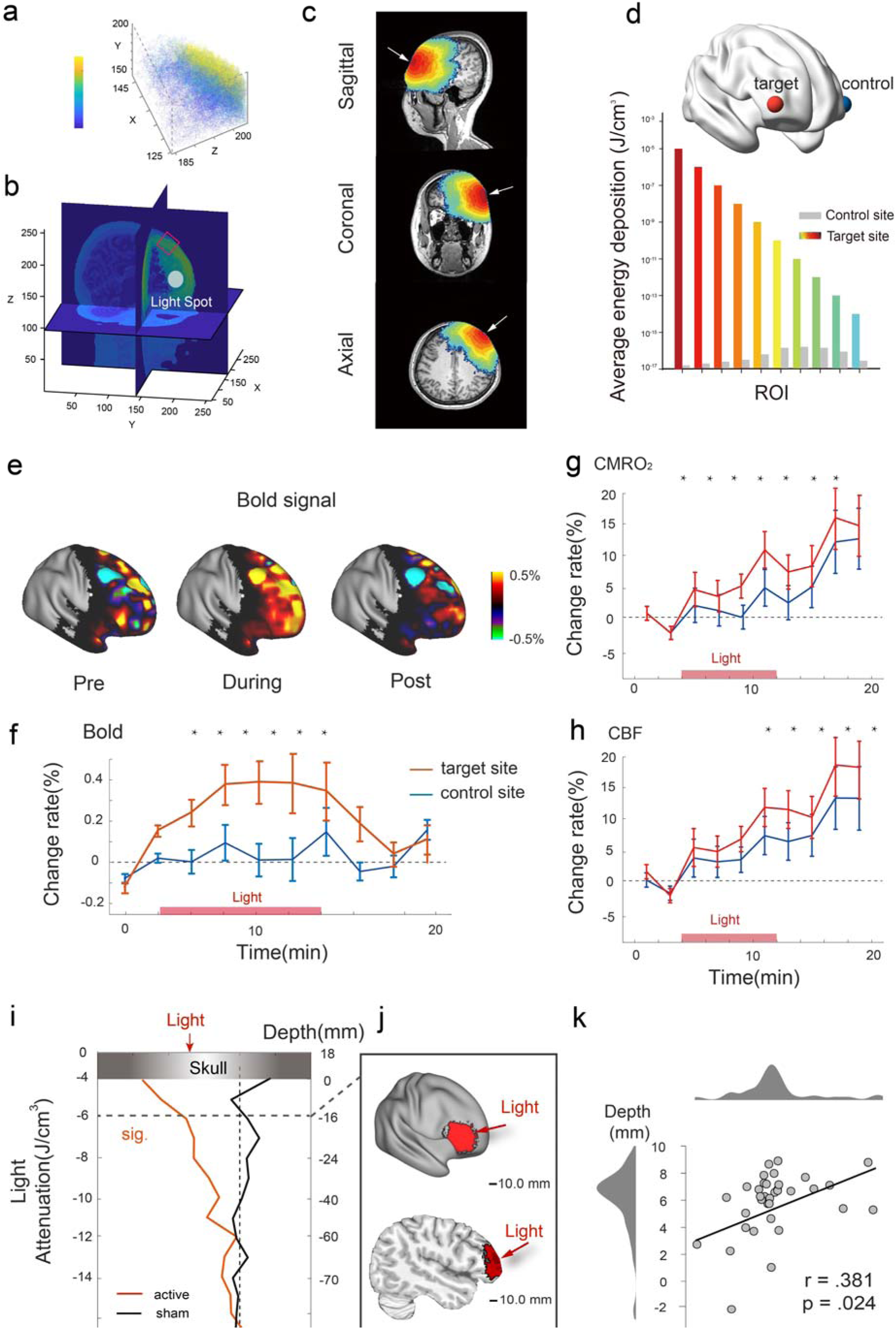
Relationship of tLS effect and light attenuation. (**a**) A close-up view of photon flux trajectories at the light spot. This illustration captures directional changes and energy attenuation at each scattering event, realistically simulating light propagation within a localized region. (**b**) The simulation is based on individualized realistic 3D head models reconstructed from structural MRI scans, incorporating major tissue classes to enable analysis of the relationship between light distribution and biological responses. The “light spot” denotes the point of light incidence. (**c**) Energy distribution is visualized on sagittal, coronal, and axial MRI slices. White arrows indicate the point of light incidence. (**d**) Average energy deposition across different ROIs (defined as the spatial domain within which photon energy attenuates by one order of magnitude). The target site, located contralateral to the control site, was selected for laser stimulation. The control site was chosen based on its anatomical and functional similarity to the target region and was not affected by incident light (power density ≈ 0). (**e**) BOLD signal activation maps at pre-, during-, and post-stimulation stages from a representative participant. Red/yellow areas indicate signal increases, whereas blue areas indicates decreases. (**f-h**) Time courses of tLS-induced changes in BOLD (**f**), CMRO_2_ (**g**), and CBF (**h**) signals. Significant increases were observed in the target region relative to baseline, compared to the control site (*p* < 0.05, two-tailed). Red lines represent the target region; blue lines represent the control region. The light pink bar denotes the stimulation period. (**i**) Light attenuation plotted as a function of depth from the scalp inward. Statistical analysis revealed a significant group-level effect of tLS under the attenuation threshold of 10^-6^, corresponding to an average effective depth of approximately 16 mm. The red line indicates the active tLS group, whereas the black line represents the sham group. (**j**) Cortical surface rendering indicating the region of significant light deposition. (**k**) A scatter plot showing the relationship between the maximum depth (10^-6^ J/cm^3^) that photons could reach and the magnitude of the tLS effect. A significant positive correlation was observed (*r* = 0.381, *p* = 0.024). Voxels with photon attenuation less than 6 dB were projected onto the cortical surface. This area, defined by the spatial extent of the tLS effect, was labeled as the tLS-ROI. 3D: three-dimensional; BOLD: blood oxygen level-dependent; CMRO_2_: cerebral metabolic rate of oxygen; CBF: cerebral blood flow; MRI: magnetic resonance imaging; ROI: region of interest; tLS: transcranial light stimulation.

To validate the physiological effects of tLS and control for potential noise or placebo responses, we implemented a sham condition. The central path of the laser defined the primary target site, while a control region that was anatomically and functionally similar but theoretically unaffected by light energy (< 10^-16^ J/cm^3^) was selected (Fig. 2d and Extended Data Fig. 2d). For subjective experience, participants wore a 10-channel stimulation array. The participants reported no awareness of specific channel activation patterns during stimulation. Their post-experiment identification accuracy (10.2% chance level) confirmed successful blinding to the stimulus locations. By analyzing the spatial features of the tLS effects and the corresponding photon energy attenuation, we quantified the light diffusion area at varying depths. Specifically, the light diffusion area (measured perpendicular to the incidence direction of the beam) initially increased with increasing depth before decreasing (Extended Data Fig. 2c). The relationship between the penetration depth and diffusion area followed a quadratic function: y=-0.705x^2^+8.767x+94.976 (*R*^2^=0.986; Extended Data Fig. 2d). Differences in brain activity between the target and control regions were examined to isolate and evaluate the specific biological effects of tLS.

### Light Simultaneously Upregulates Hemodynamic and Metabolic Activity

Previous studies have shown that tLS of the prefrontal cortex can upregulate mitochondrial CCO levels, increase HbO levels, and enhance CBF (Salehpour et al., 2018; Uozumi et al., 2010). Therefore, we hypothesized that tLS would simultaneously increase CBF and modulate both CMRO□ and BOLD signals. Given that the BOLD signal represents a complex interplay between CBF, CMRO□, and neurovascular coupling (NVC), we utilized a dual-echo sequence to acquire the BOLD and CBF signals simultaneously. This approach allowed us to computationally dissociate CMRO□ from the combined BOLD signal and CBF data, an essential step for accurately characterizing neuronal metabolic responses to tLS. To further investigate the immediate (online) and delayed (offline) effects of tLS on hemodynamic and metabolic parameters, we employed a protocol divided into three sequential phases within the MRI scanner (Fig. 1d). The session began with a 4-min baseline period (“pre-tLS”) without stimulation, followed by 8 min of light stimulation at 250 mW/cm^2^ (“during-tLS”), and concluded with an 8-min post-stimulation period (“post-tLS”) with the light turned off. As shown in Fig. 2e, the BOLD signal intensity markedly increased in the stimulated cortical region during the light exposure period relative to both pre- and post-stimulation conditions. This effect was spatially confined, with minimal changes observed in adjacent areas. The BOLD signal increased in the stimulated cortical region (> 10^-14^ J/cm^3^) during the light exposure period relative to both pre- and post-tLS (Fig. 2e). Quantitative analysis of the change in BOLD signal over time (Fig. 2 f) revealed a sustained increase in the target site beginning shortly after light onset, peaking at approximately 10–12 min, and gradually returning to baseline after stimulation ceased. In contrast, the control site exhibited no significant signal fluctuation during the same period. Two-way repeated-measures analysis of variance (ANOVA) confirmed a significant interaction between site (target vs. control) and time (*p* < 0.05), with post hoc comparisons indicating significant differences at multiple time points during stimulation.

We quantified the percentage changes in CMRO□ and CBF from baseline to represent the tLS-induced hemodynamic response. We observed significant increases in CMRO□ within the active stimulation sites from 0 to 14 min following light onset (*p* < 0.050, one-tailed; Fig. 2g), indicating sustained metabolic upregulation extending up to 6 min post-stimulation. Similarly, local CBF within the tLS-defined ROI (based on energy deposition >10^-6^ J/cm³) was significantly elevated compared to that in control regions after stimulation (*p* < 0.050, one-tailed; Fig. 2h). Notably, no significant differences were observed in baseline CBF or CMRO□ between the target and control sites (*p* > 0.050), confirming that the observed effects were attributable to tLS. These findings provide evidence that tLS induces immediate and sustained increases in local oxygen metabolism and cerebral perfusion. Importantly, this study is among the first to demonstrate simultaneous and persistent changes in CBF and CMRO□. Eight minutes of tLS not only produced an immediate increase in local CMRO□ but also maintained elevated metabolic activity in the post-stimulation period.

### Cortical Activation Patterns Correlate with Light Penetration Depth

Given the higher spatial resolution of BOLD signals, we investigated the relationship between BOLD signal changes and photon energy deposition to assess the resulting biological responses. To delineate the spatial extent of tLS within photon-attenuated ROIs, where light energy can reliably evoke biological responses, we statistically compared the stimulation effects across two independent dimensions: anatomical (target vs. control ROIs) and experimental (active vs. sham). During the active session, attenuated laser light induced significant BOLD signal changes within the target region (*p* < 0.010, two-tailed), both in comparison with the control site and the corresponding region during the sham session. We observed significant BOLD signal increases in regions receiving photon energy greater than 10^-6^ J/cm³. The measured photon distribution corresponded to a mean gray matter (GM) penetration depth of 16.0 ± 0.2 mm (standard deviation [S.D.] ± 1.2), indicating effective reach to superficial cortical layers. We further mapped the diffusion extent of −6 dB light within brain tissue. Our findings revealed that a stimulation beam with a diameter of approximately 31.1 ± 3.7 mm (S.D. ± 1.2) was sufficient to elicit measurable biological responses. Thus, this “illuminated region” (Fig. 2j) is defined as the cortical projection area within which significant tLS effects were observed; it was called tLS-ROI. Next, we examined how individual differences in the maximum depth of the tLS-ROI were related to the magnitude of BOLD signal responses within this ROI (Fig. 2k). The results revealed a positive correlation between the maximum photon penetration depth and the strength of the tLS-induced BOLD signal response (Pearson *r* = 0.381, *p* = 0.024, two-tailed). This finding suggests that efficient light transmission to the brain is associated with stronger biological responses. Our simulations show that the intracranial light field is tightly confined, with only a small fraction of light reaching the cortical surface. Nevertheless, even minimal photon absorption by brain tissue was sufficient to induce an immediate and sustained increase in the BOLD signal, confirming the sensitivity and efficacy of tLS in modulating cortical activity.

### tLS Shifts E/I Balance Toward Inhibition in Stimulated Regions

We recorded resting-state, eyes-closed scalp EEG using a 64-electrode cap and extracted source-level EEG data from the tLS-ROI, defined by simulated energy deposition exceeding 10^-6^ J/cm³ based on tLS-fMRI-ASL data (Fig. 3a). Owing to interference from the high-density EEG cap during laser transmission, the protocol was adjusted to exclude simultaneous EEG acquisition during tLS. Instead, using individualized anatomical models, we applied forward modeling and inverse solutions to reconstruct the neural activity within the tLS-ROI, thereby estimating the cortical electrical responses to intracranial photons. To strengthen the experimental control beyond the tLS-MRI setup, we introduced an active control group by changing the stimulation wavelength from the NIR-II range to the NIR-I range (see Methods for details). We first analyzed the EEG source time series using a fast Fourier transform and plotted the power spectrum in log-log space between 2 and 40 Hz (Fig. 3b). Laser stimulation led to a significant increase in theta band power (*p* = 0.032, *t* = 2.284, Cohen’s *d* = 0.666). In contrast, no significant theta band change was observed in the sham session (*p* = 0.296, *t* = 1.082, BF□□ = 0.263). Next, we explored the mechanism by which light stimulation modulated cortical excitability. Using the Fitting Oscillations and One-Over-F (FOOOF) algorithm, we decomposed the power spectrum into periodic and aperiodic components. The aperiodic slope served as an indirect marker of the E/I balance, with steeper slopes reflecting an increased inhibitory tone (Fig. 3c). We first calculated the changes in functional E/I (fE/I) for each condition. Participants who received NIR-II tLS exhibited significantly reduced fE/I after stimulation compared to those in the sham group (mean = −0.168, standard error of the mean [SEM] = 0.077; *p* = 0.038, *t* = −2.200,

**Fig. 3.**
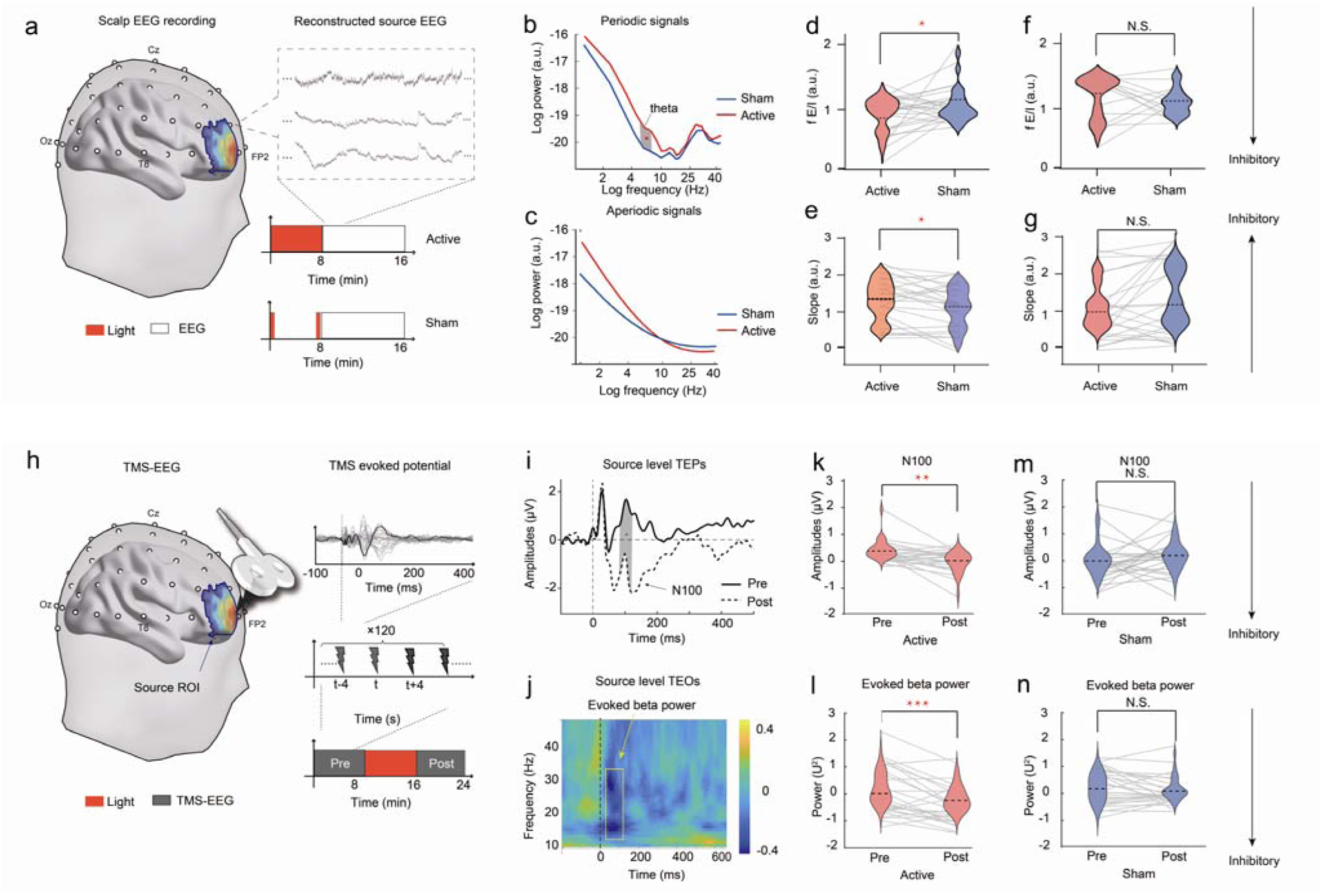
Effects of tLS on EEG activity and TMS–EEG response. (**a**) Schematic representation of the EEG electrode placement (white dots) on the scalp, with a heatmap illustrating the photon energy absorption spectrum within the tLS-ROI. The inset displays sample EEG data at source levels. The entire experiment consisted of an 8-min stimulation period (indicated in red) followed by an 8-min non-stimulation period (indicated in gray). As the stimulation site overlapped with the electrode position, we recorded EEG only after the stimulation period. (**b**) Log-transformed power spectral density (PSD, a.u.) of rhythmic neural oscillations across frequency bands under active (red) and sham (blue) stimulation conditions in the NIR-II spectrum. (**c**) Log-transformed PSD (a.u.) of aperiodic neural oscillations across frequency bands under active (red) and sham (blue) stimulation conditions in the NIR-II spectrum. (**d**) Violin plot shows the distribution of the fE/I (a.u.) under active and sham conditions in the NIR-II spectrum. Lower values indicate increased inhibitory activity. Lines represent the fE/I scores of individual participants, depicting the change in the fE/I ratio between the active stimulation (red) and sham (blue) conditions. A significant difference is denoted by an asterisk (\**p* < 0.05). Black lines represent mean scores ±SEM across each group. (**e**) Violin plot shows the slope distribution of the aperiodic exponent (a.u.) under active and sham conditions in the NIR-II spectrum. Higher values indicate increased inhibitory shifts of balance. The lines demonstrate the shift in the aperiodic exponent following active stimulation (red) compared to sham (blue). (**f**) Violin plot shows the distribution of the fE/I (a.u.) under active and sham conditions in the control group. (**g**) Violin plot shows the slope distribution of the aperiodic exponent (a.u.) under active and sham conditions in the control group. **(h)** TMS–EEG protocol for cortical response of tLS as well as pre- and post-tLS; 120 pulses of magnetic stimulation at 100% threshold intensity were delivered to the tLS-ROI at 4-s intervals. A 62-channel EEG was collected, and source localization was performed to determine the time series within the tLS-ROI. For Tth timepoint, the magnetic pulse induced an EEG response (black line) as well as responses at other time points (gray lines). **(i)** Source-level TEPs at pre- and post-tLS sessions. Grand average of EEG responses from 100 ms pre- to 500 ms post-transcranial magnetic pulse at source ROI, recorded during pre-tLS (red line) and post-tLS sessions (black line). The N100 component shows a more negative trend between 80 and 130 ms. (**k)** In active session, the mixed-model analysis of variance revealed a significant interaction effect of condition (active and sham) and sessions (pre- and post-tLS) on the change in the N100 component (*F*_2,82_ = 23.200, *p* < 0.001, two-tailed). Post hoc tests indicate that tLS increased the amplitude of the N100 component (more negative) in the tLS-ROI (*t*_41_ = 8.565, *p* < 0.001, two-tailed). **(m)** No statistically significant changes of TEPs were observed in the sham session. **(j)** Difference in TEOs at the source level between pre- and post-tLS sessions. A time-frequency map of the evoked oscillation power (post-tLS minus pre-tLS) shows a significant decrease within the frequency range of 14–30 Hz from 60 ms to 130 ms. **(i)** The mixed-model analysis of variance revealed a significant interaction effect of condition (active and sham) and sessions (pre- and post-tLS) on the change in TEOs (*p* < 0.001, two-tailed). Post hoc tests indicate that tLS reduced beta power in the tLS-ROI region (*t*_41_ = 8.565, *p* < 0.001, two-tailed). **(n)** No statistically significant changes of TEOs were observed in the sham session. EEG, electroencephalogram; fE/I: functional excitation/inhibition; NIR: near infrared; ROI: region of interest; SEM; standard error of the mean; tLS: transcranial light stimulation; TMS: transcranial magnetic stimulation; TEP: TMS-evoked potential; TEOs: TMS-evoked oscillations.

Cohen’s *d* = −0.449; Fig. 3d), indicative of increased inhibition. To further validate the inhibitory effects of the aperiodic slope, linear fits of the power spectrum in log-log space revealed significantly steeper slopes following light stimulation (mean = 0.169, SEM = 0.068; *p* = 0.021, *t* = 2.480, Cohen’s *d* = 0.506), suggesting enhanced inhibition (Fig. 3e). In contrast, the aperiodic slope showed no significant change under the sham condition (*p* = 0.520, *t* = 0.656, BF□□ < 0.294; Fig. 3g). Conversely, participants in the NIR-I group showed no significant change in theta band power, fE/I, and slope (*p* > 0.231, *t* < ±1.041, BF□□ < 0.306; Fig. 3e,f). Therefore, these findings suggest that NIR-II tLS modulates the local E/I balance by enhancing inhibitory activity. This is reflected in both the increased aperiodic spectral slopes and reduced fE/I ratios, providing converging evidence of light-induced inhibitory perturbations in local cortical activities.

### TMS-EEG Evidence of tLS-Induced GABAergic Inhibition in Human Cortex

We used a figure-of-eight TMS coil (Fig. 3h), positioned at the light stimulation target site, to deliver 120 single magnetic pulses (4-s inter-pulse interval) across two 8-min sessions—one before and one after tLS. TMS–EEG is a non-invasive method for assessing cortical excitability. This protocol was designed to investigate how tLS modulates cortical excitability by analyzing the TMS-evoked potentials (TEPs) within the tLS-ROI. TEP amplitudes, which reflect cortical excitability, were monitored using electrodes placed near the tLS-ROI. Source-localized event-related potentials were observed in both pre- and post-tLS sessions. Post-tLS TEPs demonstrated a significant reduction in amplitude during the 80–110-ms time window, corresponding to the N100 component (Fig. 3i). To examine this effect in greater detail, we conducted a two-way mixed ANOVA on the N100 amplitudes, with condition (active vs. sham) and session (pre- vs. post-tLS) as factors. A significant condition-by-session interaction was found (*F*_4,116_ = 3.471, *p* = 0.037, η^2^ = 0.314; Fig. 3k, m). Specifically, the N100 amplitude was significantly reduced following active tLS (*t*_25_ = −2.266, *p* = 0.031, Cohen’s *d* = −0.414; Fig. 3k), whereas no significant change was observed under the sham condition (*ps* > 0.250, BF_10_ < 0.333; Fig. 3m). Previous pharmacological studies have shown that the N100 component is modulated by GABAergic receptor activity, establishing it as a reliable marker of GABAergic intracortical inhibition (Premoli et al., 2014; Du et al., 2018). Thus, the tLS-induced reduction in N100 amplitude likely reflects a localized enhancement of cortical inhibition. In addition to TEP analysis, we computed the change in TMS-evoked oscillations (TEOs) by subtracting pre-tLS from post-tLS time–frequency representations. Paired *t*-tests with false discovery rate correction (*p* < 0.050) revealed a significant reduction in the beta band (14–30 Hz) power during the 60–130-ms window (Fig. 3j). A two-way mixed ANOVA confirmed a significant interaction between condition and session (*F*_1,22_ = 1.170, *p* = 0.0291, η^2^ =0.110). Post hoc *t*-tests showed a significant beta power reduction after tLS under the active condition (*t*_22_ = 2.313, *p* = 0.030, Cohen’s *d* = 0.463; Fig. 3l), whereas no significant changes were found under the sham condition (*p* > 0.351; Fig. 3n). Moreover, the tLS-induced change in N100 amplitude correlated significantly with the change in evoked beta power (*r* = −0.430, *p* = 0.023, two-tailed), suggesting a shared inhibitory mechanism. The observed reduction in TMS-evoked beta oscillations is consistent with enhanced cortical inhibition (Rocchi et al., 2018). Hence, both TEP and TEO analysis findings support the conclusion that NIR-II tLS induces a measurable increase in cortical inhibitory activity within the tLS-ROI.

### Modeling tLS-Induced Neurovascular Coupling: GABAergic Inhibition Drives NO-Mediated Vasodilation

To further explore the effects of tLS, we adopted a quantitative NVC model framework (Sten et al., 2023; Fig. 4a). Within the defined tLS-ROI, we extracted key physiologic responses (CMRO□, CBF, and BOLD signal) as model observables. As shown in Fig. 4b, the model was trained using experimental data collected from both the target and control sites, including BOLD signals (Fig. 2f), CBF responses (Fig. 2h), and CMRO□ changes (Fig. 2g). Informed by the findings of Experiments 2 and 3, tLS was implemented in the model as an inhibitory input (µ□). Parameter estimation was performed to achieve a good fit with the observed data (Jlsq = 54.64, cut-off: χ² for 60 data points = 148.78) using an objective function minimization approach to optimize the NVC parameters (see Methods section for details). Using the calibrated model, we simulated the temporal dynamics of neuronal activity in GABAergic interneurons and pyramidal neurons (Pyrs) under light stimulation at the target and control sites (Fig. 4c). The model also predicted upstream vasoactive signals mediated by partial oxygen pressure and NO (Fig. 4d), as well as downstream vascular responses, including CBF and cerebral blood volume (CBV) (Fig. 4e). The model outputs revealed clear multilevel distinctions between target and control regions along the neurovascular cascade. In the neural domain (Fig. 4c), tLS triggered a pronounced increase in GIN activity and concurrent suppression of Pyr activity in the target region, and these patterns were absent in the control site, indicating a shift toward inhibitory dominance. In the vasoactive domain (Fig. 4d), vascular smooth muscle-derived NO (NOvsm) level rose sharply in the target region during stimulation, whereas vascular smooth muscle-derived prostaglandin E□ (PGE□vsm) level showed a more gradual increase in the control region, suggesting distinct pathways of vascular regulation. These upstream modulations were reflected in vascular outcomes (Fig. 4e), where both CBF and CBV exhibited significantly greater increases in the target region, consistent with enhanced NO-mediated vasodilation. Collectively, the model-based findings delineate a mechanistic pathway through which tLS promotes inhibitory neural activity and drives NO-dependent vascular responses, specifically within the tLS-ROI.

**Fig. 4.**
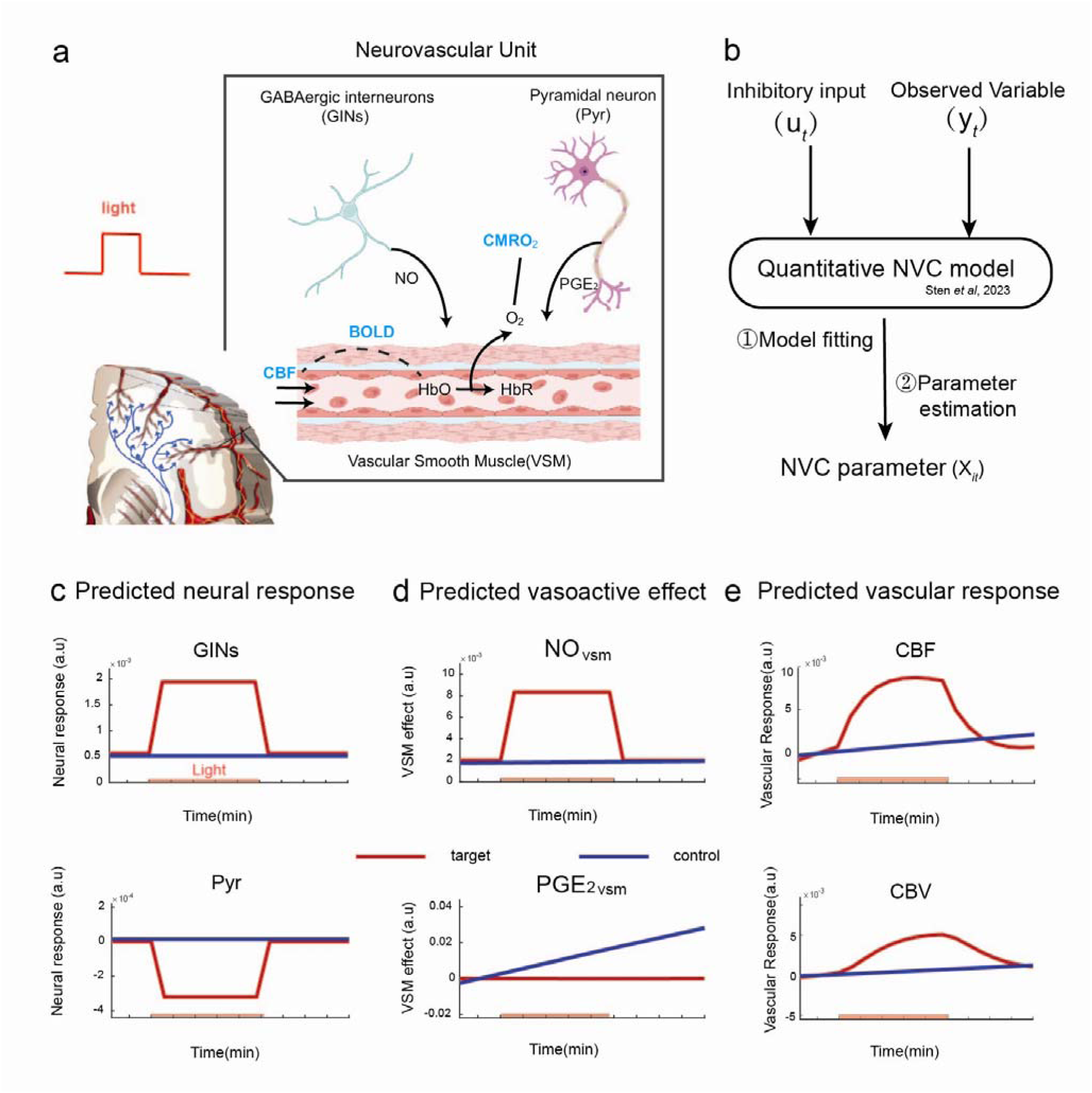
Neurovascular unit (NVU) model estimation and prediction of response. (**a**) Schematic of the NVU, highlighting GABAergic interneurons (GINs), pyramidal neurons (Pyr), vascular smooth muscle (VSM), and key biomarkers (CMROLJ: cerebral metabolic rate of oxygen; NO: nitric oxide; BOLD: blood-oxygen-level-dependent signal; CBF: cerebral blood flow; HbO/HbR: oxygenated/deoxygenated hemoglobin). (**b**) Quantitative neurovascular coupling (NVC) model (Sten et al., 2023) framework: inhibitory inputs (µLJ) and respective measurement observables (yLJ) drive model fitting and parameter estimation, yielding NVC parameters (XLJ). In **c-e**: Prediction of response under light stimulation for two sites (target sites vs. control sites). The stimulation duration (8 min) is depicted as the red bar in the bottom portion of each graph. (**c**) Neural activity of GINs and Pyr over time. (**d**) Vasoactive effects of partial oxygen pressure (POE_2_)- and NO-induced changes. (**e**) Vascular responses of CBF and CBV (cerebral blood volume) over time.

## Discussion

This study offers further insights into the mechanism by which tLS modulates brain metabolism and excitability, highlighting its potential as a neuromodulation tool that targets the NVU. We combined neuroimaging acquisition and analysis with light stimulation and employed voxel-wise mapping of hemodynamic responses along with photon transport simulations to lay the foundation for imaging-guided, standardized, and personalized stimulation protocols. Furthermore, by integrating multimodal data from EEG, TMS, ASL, and fMRI with computational modeling, we elucidated the potential metabolic pathways and signaling cascades involved in the modulatory effects of tLS, thereby advancing our understanding of light-induced neurovascular interactions.

### Biological Effects of Light Propagation in Brain Tissue

A central challenge in non-invasive brain stimulation is the effective use of external physical fields, such as electrical, magnetic, acoustic, or optical stimuli, to modulate brain activity without breaching the skull. Among these, light-based methods such as optogenetics and 40 Hz flicker stimulation utilize photons (Murdock et al., 2024), including those in the NIR spectrum. Although NIR light can penetrate the skull and affect neural activity, its direct impact on the brain tissue has not been comprehensively quantified or excluded, raising concerns regarding potential off-target effects (Dolgin et al., 2024). In response, recent optogenetic studies have begun to focus on the intrinsic interaction between light and tissues (Emiliani et al., 2022; Owen et al., 2019); moreover, inconsistencies in flicker stimulation research are increasingly attributed to variability in light. Photon variability was found to depend on parameters such as irradiance, wavelength, pulse mode, incident angle and conversion (Chen et al., 2018). Given that the biological effects of tLS depend on the absorption of light by chromophores, selecting appropriate stimulation parameters is essential to ensure therapeutic efficacy. In this regard, an accurate simulation of the experimental light source is particularly important because it determines which chromophores are likely to be activated. However, owing to inter-individual anatomical differences, photons undergo scattering and absorption by the scalp, skull, meninges, and cerebrospinal fluid (CSF), resulting in variability in intracranial light distribution (Yuan et al., 2020). Our study addresses this gap by investigating how NIR light propagates through the living human brain, characterizing its variability and revealing how it modulates blood oxygenation, metabolism, and neural excitability. Using resting-state fMRI and MC simulations, we demonstrated that NIR light can efficiently penetrate the skin and skull, with light attenuation in the brain approximating exponential decay, which is consistent with the Beer–Lambert law. According to the Grotthuss–Draper law, light absorption is a prerequisite for photochemical reactions; hence, photons must carry sufficient energy to elicit neural effects. This study marks a preliminary effort to map the spatial extent of photostimulation using BOLD signal resolution and identifies 10^-6^ J/cm³ as a possible minimal absorption dose capable of triggering biological effects. Nonetheless, considering the biphasic Arndt–Schulz law, which posits that low and high doses may induce opposite outcomes, caution is warranted when generalizing this conclusion. Further non-invasive validation of our findings in the human brain is necessary.

### Neural Excitability Modulation by tLS

We further explored the impact of tLS on cortical excitability using EEG and TMS–EEG . Previous studies have reported tLS-induced increases in alpha, beta, and gamma power (Dole et al., 2023). In this study, EEG source localization revealed that regions receiving a sufficient light dosage exhibited more negative 1/f slopes in the aperiodic EEG spectrum, indicating increased inhibition. Furthermore, TMS pulses applied to the prefrontal cortex post-tLS revealed significant reductions in the N100 component and beta band power, consistent with a shift toward an inhibitory tone. Computational modeling confirms that light stimulation increases GABAergic neuron activity while decreasing excitatory neuron activity. These results align with the observations of Cayce et al. (2011) in animal studies, demonstrating that invasive NIR light induces suppression, rather than excitation, of cortical neuron activity.

These results indicate that inhibitory neurons in the cortex appear to be more sensitive to transcranial light stimulation. This suggests that near-infrared light may selectively activate the NVU. Despite the ubiquitous distribution of mitochondria across various cell types, the differential photobiological responses observed between excitatory and inhibitory neurons might arise from their distinct spatial hierarchical organization, leading to variations in photon energy absorption. While the current study focused on excitatory and inhibitory alterations in the prefrontal region, future research should investigate whether this phenomenon exhibits region-specific characteristics. This could be achieved by employing TMS-evoked motor potentials to assess excitability in areas such as the primary motor cortex, particularly when using hair-insensitive light delivery systems. Future studies should also explore this specificty across multiple spatial and temporal scales, ranging from molecular and cellular interactions to large-scale brain networks, to validate and further elucidate the underlying mechanisms.

### Beyond Neurovascular Coupling: tLS Induces Protective Metabolic Over-Supply

Using a computational NVC model, we demonstrated a mechanistic pathway in which light stimulation enhances GIN activity and NO production, resulting in elevated BOLD signal, CBF, and CMRO□ responses in fMRI-ASL data. Our findings support the idea that tLS upregulates metabolism and blood flow, potentially via NO release and associated vasodilation (Hamblin, 2016). These effects may stem from the dissociation of inhibitory NO from CCO (Karu et al., 2008), which triggers local vasodilation and enhances metabolic support. Interestingly, we observed a decoupling of neural activity and hemodynamic responses under tLS: while excitability decreased, CBF and metabolic supply increased. This mismatch, which exceeds the actual metabolic demand, may provide neuroprotective support to vulnerable tissues. It raises the possibility that tLS may support energy availability while attenuating excitability, potentially aiding in the preservation of homeostatic balance. This effect could be particularly beneficial in conditions such as ischemic stroke or hypoxia, where oxygen delivery is impaired.

As shown in our results in Fig. 4d, tLS may induce vasodilation by generating NOvsm. Considering the upregulation of ATP by tLS, this effect can be explained in another way: tLS may also promote vasodilation by enhancing ATP production in VSMCs. Previous studies have demonstrated a significant increase in mitochondrial oxygen consumption and ATP synthesis in human endothelial cells (Wang et al., 2024).

However, our current model simplifies the complexity of the NVU, particularly in its representation of interneuron–Pyr dynamics. It omits key players, such as astrocytes, endothelial cells, and CSF dynamics, all of which contribute to neurovascular regulation. Future models should incorporate these components to understand better the systemic impacts of tLS. Similarly, our study focused on electrical and metabolic correlates, but future research using magnetic resonance spectroscopy and related techniques may uncover the involvement of additional neurotransmitters and signaling pathways.

### Long-term Effects and Future Clinical Translation

Future clinical applications depend on mechanistic insights into both the neuroprotective potential of light and its indirect effects on non-illuminated deep brain structures (Sinha et al., 2020, Hu et al., 2023). We demonstrated the feasibility of simultaneous fMRI during tLS as an in vivo method for real-time tracking of brain responses and explored the short-term direct effects of tLS. However, research (Johnstone et al., 2016; Mitrofanis, 2019) suggests that photobiomodulation induces both immediate direct effects on the NVU and prolonged secondary effects, which involve intricate regulatory processes. Thus, the long-term neuroplastic effects induced by tLS require further systematic investigation. Especially, investigating the state-specific effects of tLS is vital for the safe and effective clinical application of this technology. It is crucial for refining stimulation protocols, guiding intervention strategies, and identifying the neural markers of responsiveness.

## Materials and Methods

### Participants

In total, 128 neurologically healthy college students were enrolled in three experiments. All participants underwent medical screening and were excluded if they had a history of traumatic brain injury, neurological or psychiatric disorders, metallic implants, incompatible medical devices, or contraindications for MRI or tLS. The exclusion criteria included pregnancy, severe claustrophobia, dermatological conditions affecting the stimulation site (e.g., lesions or hypersensitivity), and metallic tattoos that could interfere with imaging. In Experiment 1, 46 participants (34 women; mean age = 23.1 years, S.D. = 1.91; range = 18.4–27.6) were recruited. In Experiment 2, 52 participants (33 women; mean age = 23.5 years, S.D. = 1.87; range = 17.2–28.7) were randomly assigned to either the NIR-II or NIR-I group. In Experiment 3, 30 participants (26 women; mean age = 24.5 years, S.D. = 2.26; range = 20.1–30.1) completed two within-participant sessions (NIR-II and NIR-I), scheduled at least 1 week apart, with the order counterbalanced across participants. Data from one participant were excluded owing to poor MRI quality. Head motion was quantified using root mean square deviation (RMSD), and participants with RMSD > 0.2 mm were excluded from further analysis. All procedures were approved by the Institutional Review Board of the Chinese Institute for Brain Research. Written informed consent was obtained from all participants, and the study was conducted in accordance with the Declaration of Helsinki.

### tLS-MRI Protocol

A neodymium-doped yttrium aluminum garnet laser was used to deliver continuous-wave light at 1064.173 nm (linewidth ±0.064 nm). Light was transmitted into the MRI environment through magnetic field-resistant optical fibers across waveguides. All optical stimulation components, including the fiber connectors, mounts, and fixtures, were constructed from nonmagnetic materials to prevent interference with the MRI static magnetic field and radiofrequency signals (Fig. 1c). Laser stimulation was synchronized with MRI data acquisition using custom control software that coordinated laser triggering with scan sequences. The fiber terminated in a beamformer mounted on a custom-designed injection-molded rubber headgear, which was divided into 10 stimulation regions and secured to the participant’s forehead for stability and positioning. VE capsules placed between stimulation sites served as fiducial markers for MRI co-registration. These markers were aligned with anatomical locations using prescanned T1-weighted structural images to accurately map the stimulation targets. The laser produced a beam spot with a diameter of 2 cm (area = 3.14 cm²), delivering 785 mW of power, corresponding to an irradiance of 250 mW/cm². The stimulation duration was 8 min, resulting in a total energy fluence of 120 J/cm² and total delivered energy of 376.8 J. The stimulation protocol consisted of a 4-min baseline MRI scan, an 8-min tLS session synchronized with ongoing scanning, and an 8-min post-stimulation MRI acquisition.

### tLS Co-registration

Owing to the limited space between the participant’s forehead and the 24-channel head coil (Siemens Ltd.), there was insufficient clearance to attach the VE markers directly to the beamformers when the participant was positioned inside the MRI scanner with the MRI-compatible laser system. To overcome this constraint, VE markers were affixed to the gaps between adjacent beamformers (Extended Data Fig. 1b). The custom headgear was contoured to match the curvature of an adult forehead, ensuring that all mounted components were positioned along an approximately spherical surface. This design optimizes both the fit and comfort of the apparatus, while maintaining precise spatial alignment. The individual beamformer positions were estimated using the coordinates of four reference VE markers (an example is shown in Extended Data Fig. 1c), based on the known geometric constraints between the beamformers and VE markers defined by the headgear design (Extended Data Fig. 1d). This approach enabled the accurate localization of the stimulation sites relative to each participant’s anatomical MRI.

We developed an algorithm to accurately estimate the positions of the 10 stimulation channels in each participant’s native T1 space. The algorithm accounts for slight deformations of the headgear that may occur because it conforms to individual differences in forehead curvature. It was assumed that all the points on the headgear were distributed on a common spherical surface. Given four noncoplanar reference points P_1_, P_2_, P_3_, and P_4_ in three-dimensional (3D) space, corresponding to identifiable VE markers, the goal was to solve for the 3D coordinates of 10 additional points lying on the same spherical surface. These points represent the beamformer positions and are constrained by the known a priori interpoint distances defined by the headgear geometry. Each estimated point *P*_i_ (*x*_i_, *y*_i_, *z*_i_) must satisfy two conditions:

1. The equation for a sphere was determined using four reference markers.
2. A set of pairwise Euclidean distance constraints between adjacent beamformer positions was defined by the fixed physical layout of the cap.

This geometrically constrained fitting approach enables precise mapping of laser incidence sites onto individual anatomical spaces, supporting the accurate co-registration of stimulation with neuroimaging data. First, the sphere parameters were determined using four reference points. A sphere is defined as follows:

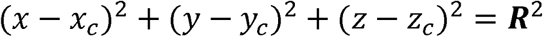

where (x_c_, y_c_, z_c_) is the sphere center and R is the radius. The distance constraints from the two reference points satisfy the following equation:

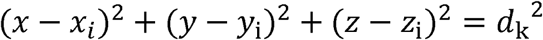

where d_k_ represents the required distances between two points.

The accuracy of the solution was validated using multiple criteria. First, all computed points were confirmed to lie on the fitted spherical surface within a numerical tolerance of less than 1×10^−6^. Second, the predefined pairwise distance constraints between adjacent channels were verified to be satisfied. Third, the reconstructed configuration was visualized using 3D plots for qualitative inspection. This approach enabled the robust estimation of 10 spatially distributed channel positions that adhered to all geometric constraints.

To achieve this, a system of nonlinear equations was formulated based on distance and surface constraints and solved iteratively to compute the 3D coordinates of each stimulation channel (Extended Data Fig. 1e). Following the initial computation, the estimated positions were reviewed via visual inspection, and minor manual adjustments were made when necessary to minimize discrepancies between the predicted and actual channel locations.

### Estimating the Illuminated Region

Transcranial light propagation was modeled using an MC simulation implemented using the MCXLAB package in MATLAB, which enables the simulation of photon migration through heterogeneous 3D media. Each participant’s head model was segmented into five distinct tissue types: the scalp, skull, CSF, GM, and white matter (WM), based on anatomical T1-weighted images processed using the CAT12 toolbox. Each voxel in the resulting atlas corresponds to one tissue category.

The optical properties of each tissue type at 1064 nm were derived from previously published literature (Extended Data Table 1). Absorption was modeled using the absorption coefficient μ_a_(1/mm), whereas scattering was defined by the scattering coefficient μ_s_. The directionality of the scattering was captured using the anisotropy factor g, where g=0 represents isotropic scattering, and g=1 indicates entirely forward-directed scattering. Brain tissues typically exhibit g≈0.9, influencing the reduced scattering coefficient μ_s_′=μ_s_(1−g). A uniform refractive index *n*=1.37 was assumed for all brain tissues, whereas air pockets within the atlas were modeled using the optical properties of air.

The light source was modeled as a top-hat beam with a flat intensity profile, and the photon injection directions were aligned along the spherical normal vectors directed toward the head center. A total of 10^9^ photons were launched per simulation, and their random interactions with the tissue boundaries, such as scattering, reflection, and absorption, were tracked in parallel using graphics processing unit acceleration. The spatially varying optical parameters necessitated continuous voxel-wise updates of the optical properties to ensure accurate calculation of the photon weight, path length, and scattering angle.

As photons propagated through the tissue, their energy was attenuated according to the Beer–Lambert law, resulting in localized volumetric energy deposition. The total source energy was normalized to 1 J, and the output was recorded as energy absorbed per unit volume (J/cm³) at a resolution of 1 × 1 × 1 mm³. These data yielded fluence maps that quantified the depth-dependent light attenuation throughout the head. The resulting volumetric light fields were visualized as color-coded intensity maps, illustrating the spatial patterns of energy absorption across tissue types.

### Neuroimaging and Preprocessing

All participants underwent MRI at the Center for Neuroimaging Sciences, Beijing University of Posts and Telecommunications, Beijing, China. Each session lasted approximately 30 min and included structural and functional imaging to acquire T1-weighted anatomical data and concurrent measurements of CBF and BOLD signals. Imaging was performed using a Siemens Magnetom Prisma 3 Tesla scanner equipped with a 24-channel transmit/receive head coil.

The scanning paradigm consisted of a 4-min baseline acquisition, an 8 min session of concurrent tLS and fMRI, and an 8-min post-stimulation acquisition. tLS was manually activated and deactivated during sessions. Throughout the scan, participants rested with their eyes closed to minimize sensory and cognitive confounding factors.

T1-weighted structural images were acquired using a magnetization-prepared rapid gradient-echo sequence (field of view = 230 mm; in-plane resolution = 256 × 256; 224 slices; slice thickness = 0.9 mm; inversion time = 1000 ms). A dual-echo EPI pCASL scan was performed for concurrent CBF and BOLD fMRI. Imaging parameters were: matrix size = 64×64, TE1/TE2 = 12/30 ms, TR = 4.4 s, rate-2 GRAPPA, 7/8 partial k-space, 32 slices with 3.5 mm thickness and 1.75 mm gap, 135 acquisitions with a scan time of 9 min 54 s. The tagging plane was positioned not specified mm inferior to the center of the imaging slab with a labeling duration of 1500 ms and PLD of 1500 ms.

Dual-echo EPI pCASL scans (and corresponding T1w images) were preprocessed using ASLPrep 0.5.0, which is based on fMRIPrep ,Nipype and uses FSL, ANTs, and software tools described below. The following internal software versions were used: Nipype 1.8.6, ANTs 2.3.3, FSL 6.0.5.

Anatomical processing was also performed within ASLPrep. T1w images were corrected for intensity non-uniformity using N4BiasFieldCorrection (ANTs), skull-stripped using antsBrainExtraction (ANTs) with OASIS30ANTs as the target template, and spatially normalized to the MNI152NLin2009cAsym templates using nonlinear registration with antsRegistration.

For ASL/BOLD data processing, the middle volume of each time series was selected as a reference and brain-extracted using a custom Nipype workflow. Susceptibility distortion correction was not applied. Head motion parameters were estimated using FSL’s MCFLIRT, and motion correction was performed separately for different volume types to avoid intensity-motion confounds. Motion parameters were then concatenated, and relative root-mean-squared deviation was recalculated. Co-registration between ASL/BOLD and T1w reference images was performed using FSL’s FLIRT with 6 degrees of freedom and a boundary-based registration cost function.

Given the relatively long TR of 4.4 s, the corresponding Nyquist frequency (1/8.8 ≈ 0.11 Hz) was close to the conventional low-frequency cutoff of 0.1 Hz, indicating that additional low-pass filtering was unnecessary. To mitigate low-frequency influences such as signal drift and to avoid the risk of removing potential tLS-induced effects due to high-pass filtering, we only applied linear detrending—a minimal and conservative preprocessing step—to the BOLD data using the AFNI command 3dTcat with the -rlt++ option, which removes linear trends while preserving the overall mean of the time series.

ASLPrep calculated CBF from the ASL data using a single-compartment general kinetic model. Prior to calculating CBF, post-labeling delay values were shifted on a slice-wise basis based on the slice timing. Fractional changes in BOLD and CBF were then computed using the 4-minute pre-tLS period as the baseline.

After computing the fractional changes in BOLD and CBF signals, voxel-wise changes in cerebral metabolic rate of oxygen consumption (CMRO□) were estimated using the Davis model:

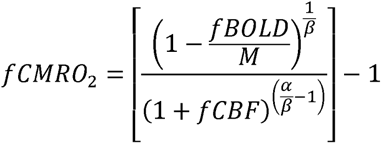

where *f*BOLD and *f*CBF represent the fractional changes in BOLD and CBF, respectively. Model parameters were set to α = 0.2 and β = 1.3. The calibration constant M was fixed at 0.07 for all participants, following prior literature.

### Illumination Uniformity of tLS

To assess the uniformity of the tLS beam profile, we followed the ANSI/NAPM IT7.228-1997 standard using a nine-point sampling method. The illuminated surface was divided into nine equal rectangular regions, and illuminance measurements were performed at the center of each region (denoted as E_1_ – E_9_) using a calibrated illuminance meter.

The average illuminance (Ea) was computed as the arithmetic mean of all the nine measurements. Illumination uniformity (U) was calculated using the following equation:

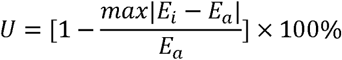

Where *max*|*E*_*i*_ - *E*_*a*_| represents the maximum absolute deviation of any measurement from the average illuminance. A uniformity value closer to 100% indicates a more homogeneous light distribution across the beam spot.

### Depth-wise Analysis of Energy Deposition

Anatomically realistic head models were generated from individual high-resolution T1-weighted MRI scans using standard segmentation into five tissue compartments: the scalp, skull, CSF, GM, and WM. Photon propagation was simulated using MCX with a spatial resolution of 1 mm³ per voxel. For each simulation, 4.2×10^9^ photons were launched from a 2-cm-diameter circular beam. The source location was set according to participant-specific experimental coordinates, and the beam direction was defined as the vector from the scalp surface to the nearest cortical gray matter voxel.

The total incident energy per simulation was normalized to 1 J, and energy deposition was quantified as absorbed energy per voxel in units of J/cm³. For each participant, two depth-resolved metrics were computed along the incident beam axis, originating from the center of the scalp projection point:

1. the energy–depth curve, defined as the voxelwise energy deposition along the beam axis; and
2. the area–depth curve, defined as the cross-sectional area (in mm²) of voxels exceeding a threshold of 10^−6^ J/cm³ in each axial plane perpendicular to the beam direction.

Inter-participant anatomical variability introduced slight differences in the spatial correspondence between depth samples and anatomical layers. To construct group-level metrics, energy and area values were averaged across participants at each depth increment. The resulting group-averaged energy–depth curve was modeled using an exponential decay function, reflecting the attenuation of light with increasing tissue depth. The area–depth curve was fitted with a second-order polynomial to capture the spatial spread of the deposited energy. Data points at extreme depths that deviated significantly from the overall trend were excluded to enhance the robustness and stability of the polynomial fit.

### EEG Recordings

Resting EEG data were recorded using the Curry 8.0 software package and a SynAmps EEG amplifier (NeuroScan Inc.). A 64-channel EEG cap with silver chloride electrodes was positioned according to the international 10-20 system, with signals referenced online to the left mastoid. Electrode impedances were maintained below 5 ㏀. EEG signals were digitized at a sampling rate of 1000 Hz. Ocular activity, including blinking and eye movements, was monitored via horizontal and vertical electrooculogram channels using external electrodes placed at the outer canthi and above and below the right eye. During the 8-min recording session, participants rested with their eyes closed to ensure a stable and alert baseline state.

### Source Analysis of EEG Signals

Source localization was performed to estimate cortical activity originating from regions targeted by tLS, based on inverse modeling of high-density EEG recordings. After the preprocessing steps, including artifact correction, bandpass filtering, and epoching, the data were imported into Brainstorm from EEGLAB for source analysis. Anatomical co-registration between EEG electrode positions and individual T1-weighted MRI anatomy was performed by identifying three fiducial landmarks: the nasion and the left and right preauricular points. EEG signals were re-referenced to the common average to satisfy the zero-net current assumption required for source estimation. The forward model was computed using the boundary element method with the default OpenMEEG settings of Brainstorm, yielding a realistic three-layer head model (scalp, skull, and brain) constructed from nested boundary surfaces. A lead field matrix was generated based on a cortical mesh with 15,000 vertices. Noise covariance matrices were computed for each participant using extended segments of resting-state data to estimate the spatial distribution of sensor-level noise. Cortical source activity was estimated using the weighted minimum norm estimation (wMNE) algorithm, with dipole orientations constrained to the cortical surface. This approach yielded dynamic current density time series at each cortical location. Time series were then extracted from tLS-ROIs, defined individually as ’*_db’ scouts in Brainstorm, based on photon energy deposition patterns derived from MC simulations. For TMS–EEG data, single-trial responses were averaged for each participant prior to source reconstruction. Source estimation was then performed on participant-averaged waveforms using the same wMNE framework.

### EEG Data Preprocessing and Analysis

EEG data were processed using EEGLAB, Brainstorm, and custom scripts in MATLAB R2020b (The MathWorks Inc.). Raw signals were downsampled to 256 Hz and band-pass-filtered between 0.1 and 50 Hz. Segments contaminated by excessive motion artifacts were manually removed through visual inspection. Noisy channels were corrected using spherical spline interpolation. Independent component analysis (ICA) was performed to remove ocular and myogenic artifacts, retaining up to four components. Artifact components were identified using ICLabel v1.6 with conservative thresholds ([0.95, 1] for muscles and [0.85, 1] for eye movement). Following artifact removal, the signals were re-referenced to the common average. On average, fewer than three blink-related components were removed per participant.

### Spectral Analysis

Power spectral density was estimated using the Welch method. Cleaned EEG data were segmented into overlapping windows, and fast Fourier transform was applied to improve spectral stability. The mean spectral power was computed for the canonical frequency bands theta (4–6 Hz), alpha (8–12 Hz), beta (14–30 Hz), and gamma (30–45 Hz) by averaging across the corresponding frequency bins.

To analyze the aperiodic component of the power spectrum, the FOOOF toolbox was applied to source-level data extracted from the tLS-ROIs. This method fits the 1/f aperiodic slope in semi-log space by iteratively removing oscillatory peaks through Gaussian subtraction. For resting-state recordings, only the 50 Hz line noise and the 8–12 Hz alpha peak were consistently identified and excluded from the final slope estimation. This procedure provided a robust characterization of non-oscillatory broadband activity within the tLS-ROI.

### Calculation of Functional E/I Balance

To estimate the fE/I balance, we adopted the method described by Bruining et al. (2020). For each EEG channel, the signal profile *S*(*t*) was constructed as the cumulative sum of the demeaned amplitude envelope *A*(*t*):

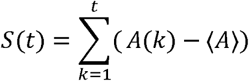

The signal was segmented into overlapping windows of 5,000 samples with 80% overlap. Within each window, amplitude normalization was performed by dividing the windowed signal by its mean amplitude, thereby reducing bias from nonstationary fluctuations. Detrending was applied to eliminate linear trends and stabilize temporal dynamics.

The normalized fluctuation function nF(t) was then computed as the standard deviation of the detrended normalized signal in each window. Long-range temporal correlations were quantified using detrended fluctuation analysis, and the fE/I balance was calculated as the Pearson correlation between the original windowed amplitude and the corresponding *nF*(*t*).

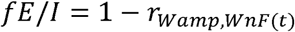

This measure reflects the degree of coupling between the signal magnitude and its temporal variability, thereby serving as a proxy for cortical E/I dynamics.

### TMS–EEG : Data Acquisition and Analysis

TMS–EEG data were acquired using a 64-channel BrainAmp DC amplifier system (BrainProducts GmbH) at a sampling rate of 2500 Hz. The electrodes followed the international 10–20 system layout, with Ag/AgCl ring electrodes mounted on an elastic EEG cap. The reference electrode was positioned at Cpz and the ground electrode at Fpz. Electrode impedances were maintained below 5 kΩ throughout the experiment to ensure signal integrity. To minimize auditory-evoked potentials, participants wore earplugs during the entire TMS session. TMS was delivered using a Magstim Rapid2 stimulator (The Magstim Co. Ltd.) equipped with a 70-mm figure-of-eight coil. Before the experimental session, the resting motor threshold (RMT) was determined for each participant according to established international guidelines. The RMT was defined as the minimum stimulation intensity required to elicit motor evoked potentials with an amplitude ≥50 µV in at least five out of 10 trials while the target muscle was at rest.

The formal TMS–EEG experiment consisted of three stages: a pre-tLS baseline, a 12-min tLS modulation phase targeting the right frontal cortex, and a post-modulation measurement phase. TMS–EEG recordings were acquired immediately before and after the tLS session. In each pre- and post-modulation stage, 120 single TMS pulses were delivered at 4-s intervals to the tLS-ROI with the stimulation intensity set at 100% of the individual’s RMT. EEG signals were continuously recorded during stimulation.

The continuous EEG data were segmented into epochs from –1000 ms to +1000 ms relative to the TMS pulse onset and downsampled to 256 Hz. To suppress TMS-induced artifacts, the data segment from –2 ms to +20 ms around the TMS pulse was removed and interpolated using the cubic interpolation function in the TMS–EEG Signal Analyzer toolbox. A fourth-order zero-phase Butterworth filter was applied, including a 48–52 Hz notch filter to suppress line noise and a 1–100 Hz band-pass filter to preserve the relevant EEG frequencies.

Artifact removal was performed using ICA. Components related to ocular and myogenic artifacts were identified based on frequency content and topographical features and removed manually. Finally, the data were re-referenced to the global average across all electrodes to prepare for source localization and further analysis. To assess cortical responses to TMS, both time-domain and time–frequency domain analyses were performed based on source-reconstructed signals within the tLS-ROI. Specifically, TEPs were computed to characterize transient voltage deflections following stimulation, whereas event-related spectral perturbation (ERSP) analysis was employed to examine frequency-specific changes in power. For the ERSP analysis, source-level data were transformed using Morlet wavelet decomposition. The time window from –200 ms to +800 ms relative to TMS onset was retained for time–frequency decomposition, and a pre-stimulus baseline from –800 ms to –400 ms was used for normalization. Baseline correction was performed to remove power-law effects in the EEG spectrum following the normalization procedure defined as follows:

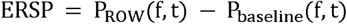

Morlet wavelets were parameterized with a center frequency of 1.5 Hz and a spectral bandwidth of 1 Hz, enabling adaptation across the relevant EEG frequency range. This configuration provided a balance between temporal and spectral resolutions suitable for capturing event-related oscillatory dynamics in the theta to beta bands.

### NVC Model

To characterize the physiological mechanisms underlying tLS effects on brain function, we employed a quantitative NVC model comprising three integrated modules: neuronal activity, vascular dynamics, and oxygen transport and BOLD signal generation. These modules were embedded within a unified mathematical framework to simulate a cascade of neural, vascular, and metabolic processes.

The neuronal module captures interactions between excitatory pyramidal cells and inhibitory interneurons, including those releasing NO, PGE□, and neuropeptide Y (NPY). These neurotransmitters modulate the vascular tone and metabolic demand. The vascular module simulates blood flow, pressure, and volume changes across arterioles, capillaries, and venules. The oxygen transport module models oxygen diffusion and hemoglobin dynamics (HbO, HbR, and HbT) and predicts the resulting BOLD signal. The model was formulated as a hybrid system of ordinary and differential algebraic equations and numerically solved using the AMICI simulation toolbox. Initial and boundary conditions were derived from empirical literature or calibrated to experimental datasets. Parameters were estimated using the MEIGO toolbox with an enhanced scatter search (eSS) algorithm to efficiently explore the high-dimensional parameter space. Fitting was performed by minimizing the negative log-likelihood function under the assumption of Gaussian noise.

The CMRO□ was modeled as a function of neuronal activity.

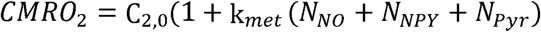

where N_NO_, N_NPY_, and N_Pyr_ denote the activity levels of NO-expressing interneurons, NPY interneurons, and Pyr, respectively, and k_met_ is a scaling parameter. Oxygen transport is governed by the compartmental exchange across the arterial, capillary, and venous compartments. Oxygen extraction occurs within the tissue space, generating the metabolic signal CMRO□.

The BOLD signal was calculated as the weighted sum of the contributions from each vascular compartment.

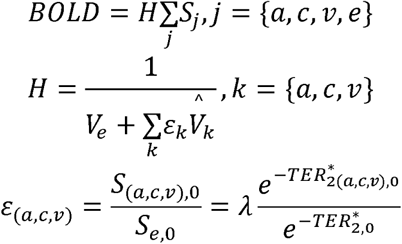

where S_j_ represents the intrinsic signal from each compartment *j*, V_e_ is the extracellular volume, V^-^_k_ is the intravascular volume, and λ = 1.15 is the spin-density.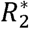 represents the transverse relaxation rate, and TE is the echo time of the fMRI sequence.

Model parameters θ were estimated by minimizing the negative log-likelihood:

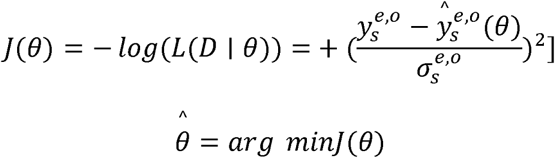

where 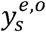 and 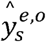 denote observed and simulated values, and 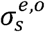 is the standard deviation of the data.

Optimal parameters θ̂ were obtained by solving 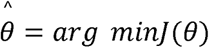

Uncertainty quantification was conducted using Markov chain MC sampling with 10^5^ samples. Posterior distributions were evaluated using a chi-squared goodness-of-fit test (*p* < 0.05). Acceptable parameter sets satisfied the following equation:

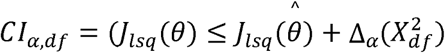

where 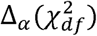 is the α quantile of the chi-squared distribution. Parameters producing implausible or unstable dynamics were excluded from further analysis.

**Extended Data Fig. 1.**
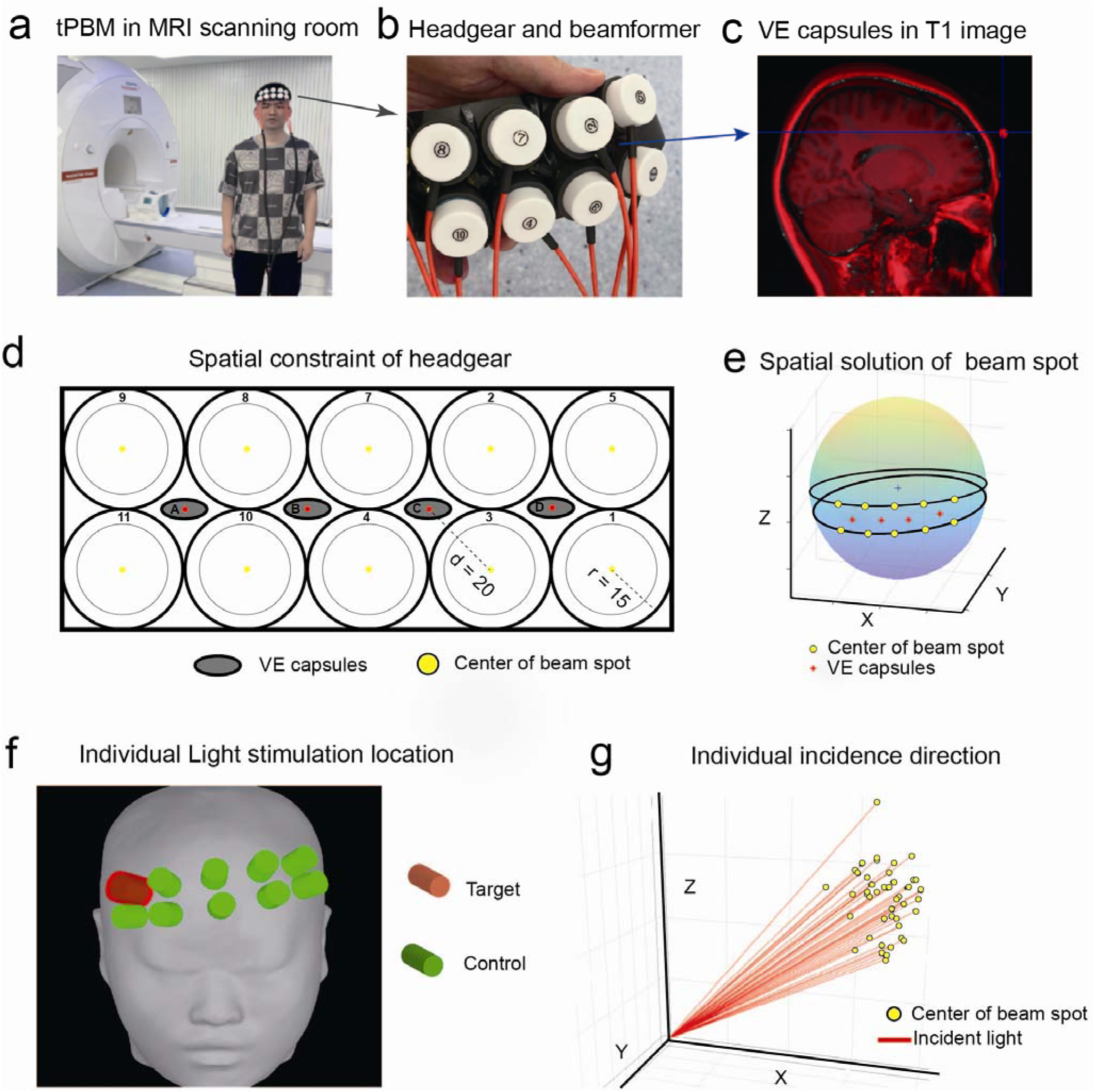
**(a)** The author himself wearing the MRI-compatible headgear and tLS setup inside the MRI scanner (Siemens Prisma 3.0T). **(b)** Close view of the multichannel tLS setup. The customized rubber headgear can accommodate 10 laser light beamformers. Because of the limited space within the MRI coil, VE markers are placed at the center of the beamformers. **(c)** By co-registering two T1 structural images (with and without spatial markers), the coordinates of the VE capsules (red dots) in the T1 image can be determined. **(d)** Schematic of the tLS headgear with spatial markers (gray ellipses) placed at the center of 10 light stimulation rings. The outer diameter of the laser light beamformer is 15 mm, and the distance from the center of the VE capsules to the center of the beam spot is 20 mm. **(e)** Using the known spatial coordinates of the four VE capsules and the spatial constraints of the headgear, the coordinates of the 10 tLS centers can be calculated at the individual level. **(f)** The upper-right position (red light column) is selected as the target area for stimulation for each participant, with the other nine green light columns serving as control areas. **(g)** The individualized incidence direction of light stimulation is shown, with yellow dots indicating the center of the beam spots. MRI: magnetic resonance imaging; tLS: transcranial light stimulation; VE, vitamin E.

**Extended Data Fig. 2.**
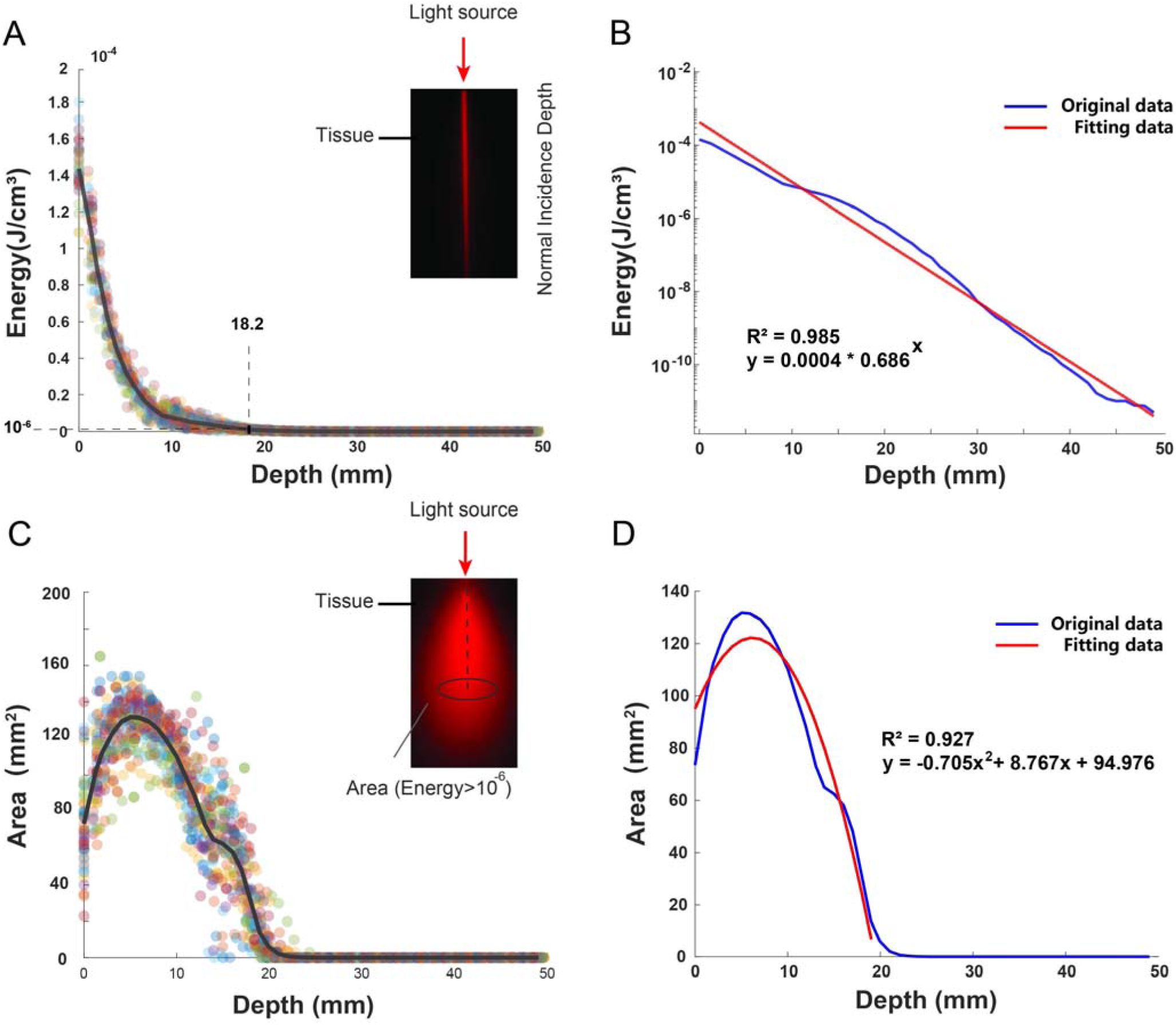
Depth-dependent intracranial light distribution under tLS. **(a)** Voxel-wise energy deposition (in J/cm³) was computed for each participant along the individualized light incident path, with depth measured from the center of the projected beam footprint on the scalp surface. Each colored dot represents one participant, and the black line represents the group average. The dashed horizontal line marks the energy threshold (10LJLJ J/cm³), and the dashed vertical line indicates the depth at which the group-averaged energy reaches this threshold (18.2 mm). *Inset*: Spatial distribution of energy deposition along the incident light path. **(b)** Semi-logarithmic plot of the group-averaged energy–depth profile fitted using an exponential decay model (*R*² = 0.985), showing the attenuation of photon energy with increasing depth. **(c)** Cross-sectional area (in mm²) was computed at each depth plane orthogonal to the incident path by summing the area of voxels where energy deposition exceeded 10LJLJ J/cm³. The depth axis is defined identically to that in (a). Colored dots represent individual participant data, and the black line denotes the group average. *Inset*: Spatial distribution of intracranial energy deposition across the depth and lateral directions aligned with the incident path. **(d)** The group-averaged area–depth profile was fitted using a second-order polynomial (*R*² = 0.927), revealing a bell-shaped pattern of lateral light spread across the depth. Data points at greater depths that deviated substantially from the overall trends were excluded to improve the stability of the fit. tLS, transcranial light stimulation.

**Extended data Table 1 Optical parameters of brain tissues.** Mus: scattering coefficient (1/mm); mua: absorption coefficient (1/mm); g: anisotropy factor; n: refractive index; GM: gray matter; WM: white matter; CSF: cerebrospinal fluid, as adopted from a previous study (Cassano et al., 2019).

